# A method for identifying spatially divergent selection in structured populations

**DOI:** 10.1101/2025.02.23.639751

**Authors:** Isabela do O, Oscar Gaggiotti, Pierre de Villemereuil, Jerome Goudet

## Abstract

Species occupy diverse, heterogeneous environments, which expose populations to spatially varied selective pressures. Populations in different environments can diverge due to local adaptation. However, neutral evolution can also drive population divergence. Thus, testing for local adaptation requires a neutral baseline for population differentiation. The classical *Q_ST_* -*F_ST_* comparison was developed for this purpose. Yet, *Q_ST_* -*F_ST_* frequently fails to account for the complexities of population structure because the theory underlying this comparison assumes that all subpopulations are equally related, resulting in inflated false positive rates in metapopulations that deviate from the island model. To address this limitation we use estimates of between- and within-population relatedness to model population structure. Using those relatedness matrices, we infer the between- and within-population ancestral additive genetic variances under a mixed-effects model. Under neutrality, these inferred variances are expected to be equal. We propose here a test to detect selection based on the comparison of these two estimates of the ancestral variance and we compare its performance with earlier solutions. We find our method is well calibrated across various population structures and has high power to detect adaptive divergence.

**Author summary:** Populations of the same species often face different environmental pressures, driving them to adapt locally. However, even in the absence of adaptation, subpopulations can diverge due to random genetic drift and limited migration. Distinguishing between adaptive evolution and random divergence is a central challenge in evolutionary biology. Traditional methods, such as *Q_ST_* –*F_ST_* comparison, assume equal relatedness among subpopulations—a simplification that rarely holds in complex real-world scenarios, leading to flawed conclusions. To overcome this limitation, we developed a novel method that incorporates genetic relatedness among subpopulations, leveraging quantitative genetic theory to estimate ancestral additive genetic variances. Our approach provides a powerful tool for testing local adaptation, reliably distinguishing adaptive divergence from drift across a variety of population structures.

## Introduction

Species are generally distributed in heterogeneous environments. Over time, subpopulations exposed to different environments diverge genetically and phenotypically due to genetic drift, natural selection, and limited gene flow. Environmentally divergent subpopulations are likely to be subjected to different selective pressures, promoting their adaptive divergence. At the same time, distant subpopulations are more isolated, leading to their neutral differentiation [Lande, 1992]. Thus, before concluding that the observed difference is adaptive, one must first reject the hypothesis of neutral divergence by establishing a theoretically justified neutral expectation [Lynch et al., 1998].

*F_ST_*, the fixation index, is a classic measure used to quantify genetic differentiation among populations. For neutral loci, the degree of differentiation between subpopulations measured by *F_ST_* depends on the demographic parameters of the population, such as the effective population size and migration rates. When selection is at play, *F_ST_* for the loci under selection will also depend on the strength and mode of selection. When selection acting on a locus is similar across environments (e.g. balancing selection) *F_ST_* will be lower at this locus compared to neutral loci. When selection varies across subpopulations (e.g. local adaptation), then the selected locus will have a larger *F_ST_* in comparison to neutral loci.

Many traits of adaptive significance, such as morphological [Dittberner et al., 2018], physiological [Côte et al., 2016] and fitness-related [de Miguel et al., 2022] phenotypes, are quantitative traits and result from the interaction of multiple genes [Fisher, 1999, Lynch et al., 1998, Pritchard and Di Rienzo, 2010]. In this context of polygenic adaptation, *F_ST_* measured for neutral loci serves as the null expectation. For these traits, the standard procedure is to compare *F_ST_* with its quantitative analogue, *Q_ST_*, which describes the proportion of additive genetic variance between subpopulations relative to the total additive genetic variance [Spitze, 1993]:

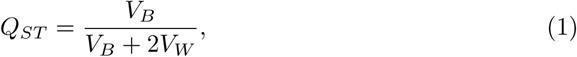

*V_B_* represents the between-population additive genetic variance, and *V_W_* is the within-population additive genetic variance. Since *Q_ST_* is based on additive genetic variances, its estimation rests on being able to use controlled breeding designs and environmental effects [Pujol et al., 2008]. Under the assumption that the investigated trait is neutral, we expect *Q_ST_* to be equal, on average, to *F_ST_*. If *Q_ST_ > F_ST_*, it suggests adaptive divergence, while *Q_ST_ < F_ST_* may imply balancing selection [Spitze, 1993, Leinonen et al., 2013].

However, *Q_ST_* estimates are subject to significant uncertainty and potential biases. First, ratio estimates, like *Q_ST_*, can be biased since the expected value of a ratio differs from the ratio of expectations [O’Hara and Merila, 2005]. Second, the number of subpopulations considered strongly affects the reliability of *Q_ST_* estimates, with fewer populations sampled increasing the variability and reducing the power of the statistic [Goudet and Büchi, 2006, Whitlock, 2008]. Because of the high variability of *Q_ST_* − *F_ST_* comparisons [Whitlock, 2008] proposed that instead of relying on a direct comparison between the two statistics, one should find where *Q_ST_* − *F_ST_* falls in the expected neutral distribution of *Q_ST_* − *F_ST_* values. Building on this, Whitlock and Guillaume [2009] developed a simulation-based approach to test whether *Q_ST_* is consistent with neutral divergence by comparing the observed *Q_ST_* − *F_ST_* to a simulated distribution of the neutral expectation. This expected neutral distribution is generated using parametric bootstrapping, drawing from a *χ*^2^ distribution following approximations by Lewontin and Krakauer [1973].

While the improvement of the *Q_ST_* − *F_ST_* method proposed by Whitlock and Guillaume [2009] reduces the false positive rate (FPR), it still assumes an isotropic population structure. For example, when considering instead a stepping-stone model, the *p*-values do not follow the expected distribution under neutrality, resulting in a non-calibrated test [Koch, 2019, de Villemereuil et al., 2022]. As highlighted in de Villemereuil et al. [2022], the main issue lies in how between-population variance (*V_B_*) is estimated in these models. Typically, *V_B_* is derived from a mixed-effects model where population-level random effects are treated as independent, assuming equal relatedness between all subpopulations. This isotropic assumption does not hold for most natural populations, which often have complex genealogical relationships and migration patterns, leading to the lack of calibration reported above.

To address this, Ovaskainen et al. [2011] proposed a model based on between- and within-population coancestry to estimate additive genetic variance accounting for non-uniform migration and drift patterns. The coancestry between two populations is the average coancestry between all pairs of individuals from those two populations, and the coancestry between pairs of individuals is the probability that randomly sampled alleles from these individuals are identical by descent (IBD). Ovaskainen et al. [2011] method extends the animal model to the metapopulation level, using coancestry to reflect genealogical relationships between subpopulations. They assume that all studied populations trace back to a common ancestral population. By accounting for population structure, their method provides a more accurate estimation of neutral expectations. Nevertheless, Ovaskainen et al. [2011] solution still relies on a metapopulation model with specific assumptions, the admixture F-model [Karhunen and Ovaskainen, 2012], which has been shown to suffer from issues in estimating coancestries [Goudet and Weir, 2023]. Besides, it is still unknown whether the test Ovaskainen *et al*. developed can be considered calibrated, given that it does not provide a null-hypothesis framework *per se*.

In this study, we introduce LogAV, a new method for testing the null hypothesis of neutral divergence by comparing the log-ratio of two estimates of the same ancestral additive genetic variance: one derived from a between-population effect and the other from a within-population effect. We then evaluate this method along with both Whitlock and Guillaume [2009]’s method and the Driftsel approach [Karhunen et al., 2013] which implements Ovaskainen et al. [2011] ideas. We assess the performance of LogAV through simulations of neutrally evolving populations, considering a range of population structures, including highly non-isotropic configurations. We found LogAV well calibrated for all tested population structures and able to detect selection when present.

### LogAV description

LogAV is designed to test whether neutral processes alone can account for the observed divergence between related subpopulations of a metapopulation. We assume that all studied subpopulations originated from a single, panmictic, ancestral population, and through a process of divergence and restricted gene flow, their genetic diversity becomes spatially structured. As shown by Ovaskainen et al. [2011], it is possible to write an inferential model to estimate the expected ancestral additive variance (*V_A_*) under neutrality. We define the ancestral additive genetic variance *V_A_* as the additive genetic variance in the quantitative trait that existed in the ancestral population. This model provides a baseline for understanding the contributions of neutral processes and selection to the current genetic makeup of each subpopulation. Our approach examines the historical processes that occurred as they diverged from their common ancestor, as well as the ongoing evolutionary process, including current migration patterns and inbreeding.

In quantitative genetics, an animal model is generally used to estimate additive genetic variance. This model partitions phenotypic variance into different genetic and environmental components. To quantify the covariance of genetic effects between individuals, this model depends on the use of an individual-level relatedness matrix. Extending this principle to structured populations requires quantifying relationships between subpopulations as well as between individuals within those subpopulations [Ovaskainen et al., 2011]. Demographic history and spatial metapopulation configuration influence the distribution of genetic variation, which affects additive genetic variance within and between subpopulations. Thus, any test of local adaptation requires that the neutral differentiation baseline used accounts for these effects.

LogAV estimates *V_A_* in two separate ways: using the between-population and the within-population coancestries. We then compare these two estimates of the additive ancestral variance. We used a method of moment estimator of coancestries [Goudet and Weir, 2023] to measure (i) kinship within subpopulations and infer ancestral variance from within-population variance, and (ii) mean kinship between subpopulations and infer ancestral variance from between-populations variance. Since the mathematical framework is constructed to reference the same coalescence tree, it accurately represents the segregation of alleles associated with the same variance. Thus, in the absence of selection, both variances should be equal.

To illustrate the concepts above, let us first consider a scenario involving two subpopulations. In this simple case, we can use *F_ST_* to characterize the population structure of our metapopulation to establish the expected relationship under neutrality between the current genetic variances and the ancestral genetic variance. Namely, the proportion of the total genetic diversity described by the diversity between subpopulations is 2*F_ST_*, and the remaining proportion, 1 − *F_ST_*, is the one within subpopulations. Thus, at the limit of low mutation rate, and under neutrality, we should expect that the between-population additive genetic variance (*V_B_*) and the within-population additive genetic variance (*V_W_*) are respectively:

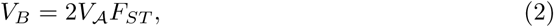

and

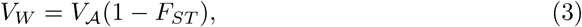

as demonstrated by Whitlock [1999], and earlier by Wright [1969].

However, as noted by others before [Weir and Hill, 2002, Karhunen and Ovaskainen, 2012, Weir and Goudet, 2017] *F_ST_* is a global summary parameter that does not capture differences in demographic histories among subpopulations under complex population structures. These differences in demography could be for example variation in population size, different branching times between subpopulations, and variation in migration rates between pairs of subpopulations. To capture the complexity of more biologically relevant population structures, we use a matrix of coancestries **Θ***^p^* following Ovaskainen et al. [2011], Goudet and Weir [2023].

**Θ***^p^* contains the expected coancestries between pairs of subpopulations relative to the ancestral population (Fig 1). The superscript “*p*” refers to it being a population-level parameter. Element Θ*^p^* describes the probability of IBD between a random pair of alleles, one coming from subpopulations *i* and the other from subpopulation *j*. Diagonal elements of **Θ***^p^* describe such probability for any pair of alleles from distinct individuals within a subpopulation. **Θ***^p^* accounts for the non-independence between subpopulations, shared evolutionary history, asymmetric migration rates, and differences in population sizes, addressing limitations of using the global summary parameter *F_ST_* in unevenly structured metapopulations [Goudet and Weir, 2023]. *F_ST_* and **Θ***^p^* are related as *F_ST_* is proportional to the average of the diagonal elements of **Θ***^p^*:

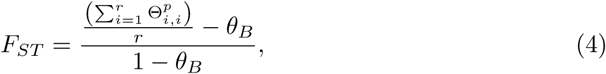

**Fig 1.**
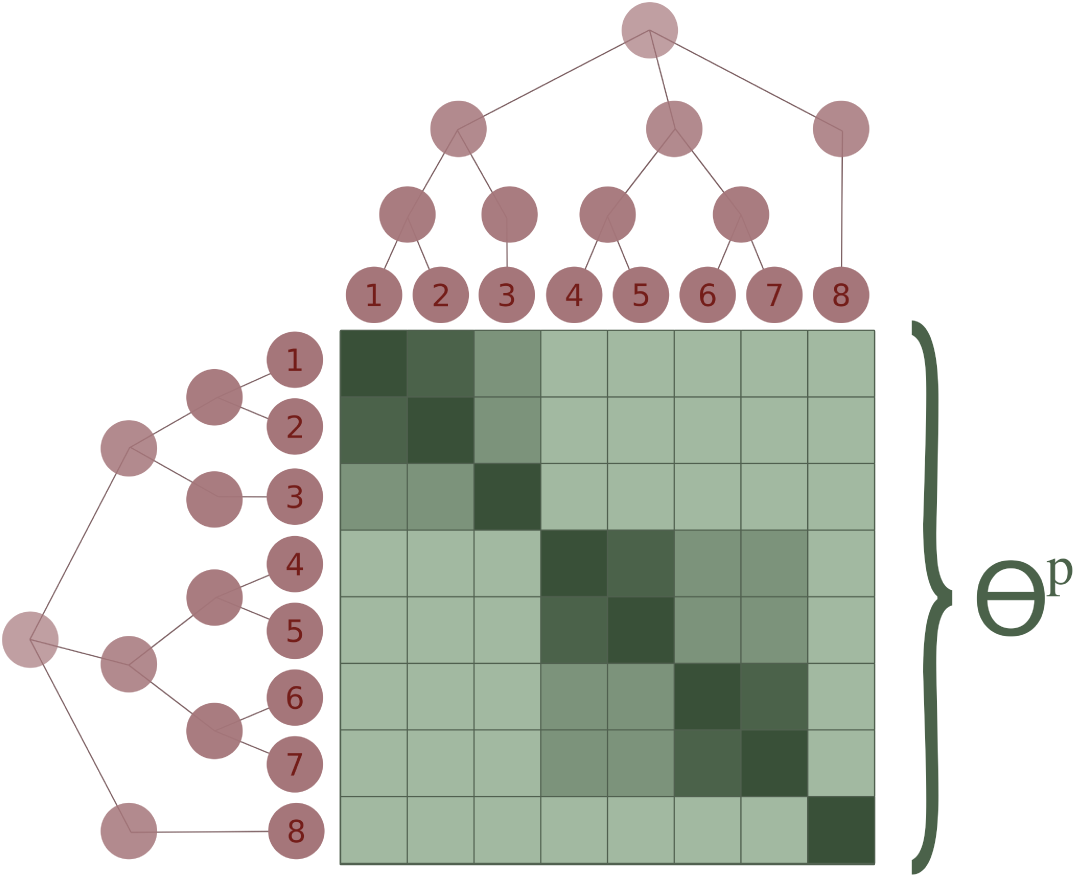
Cartoon description of **Θ***^p^*. Given *r* subpopulations, **Θ***^p^* will be a *r* × *r* matrix. In this figure **Θ***^p^* is represented in different shades of green describing the average level of coancestry between random alleles sampled from different pairs of subpopulations, with the diagonal being the average of random pairs of alleles from distinct individuals in the same subpopulation. Darker shades in the cells within the matrix represent higher coancestry between the pair of subpopulations being compared. **Θ***^p^* reflects the past and present demographic metapopulations’ histories and connectivity.

where 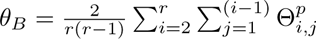 is the average of the off-diagonal elements of **Θ***^p^*.

While **Θ***^p^* describes average co-ancestries across and within subpopulations, the **M** matrix [Fig. 2, Ovaskainen et al., 2011] describes relatedness between pairs of individuals. As it is general practice, we assume no cross-population breeding was performed, and thus that **M** is a block diagonal matrix [this assumption can be lifted and cross-population breeding accounted for in **M**, see Ovaskainen et al., 2011], *i.e.* relatedness for pairs of individuals from different subpopulations are already defined with **Θ***^p^*, so they are set to zero in **M**. The blocks in **M** describe the relatedness between all pairs of individuals within the subpopulations, relative to the average relatedness within each subpopulation, and discounting relatedness already accounted for in **Θ***^p^*. More precisely, **M** contains, within each subpopulation block *x*, the relatedness for each pair of individuals in that subpopulation multiplied by 1 − Θ*^p^_x,x_*.

**Fig 2.**
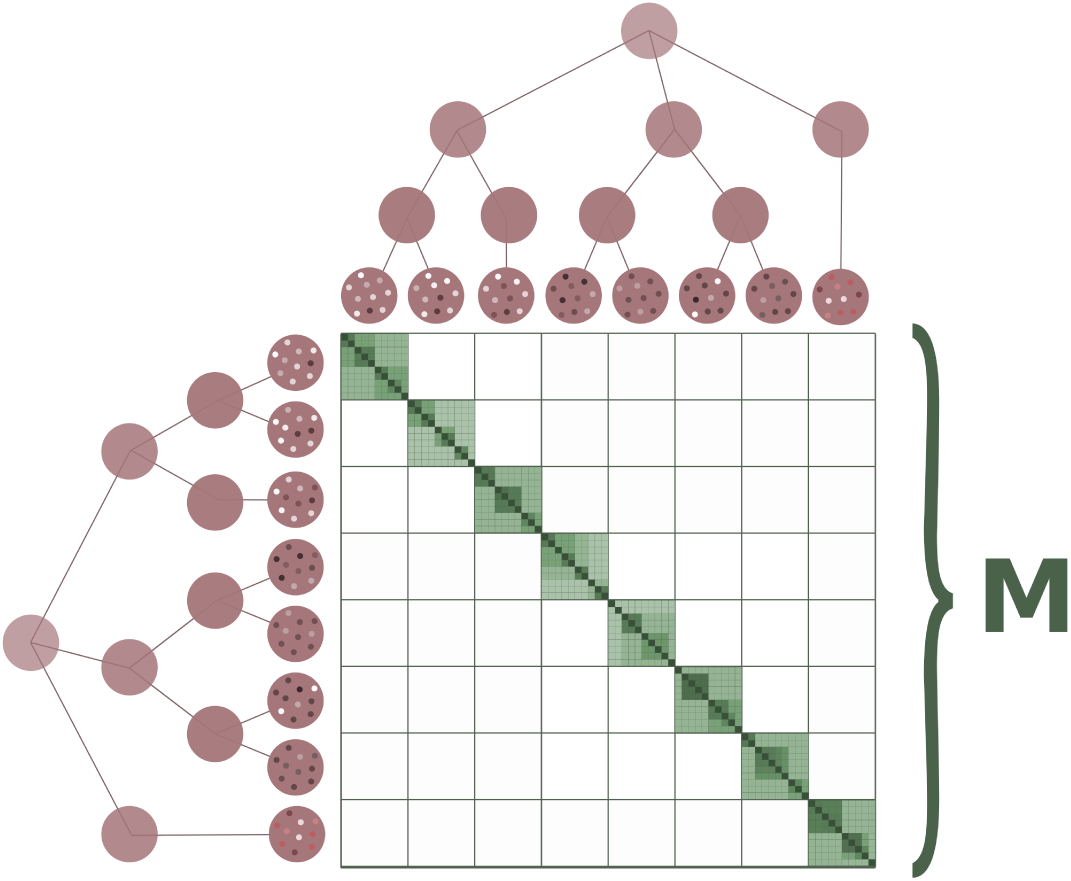
Cartoon description of *n_T_* × *n_T_* matrix **M**. Similar to Fig 1, darker shades describe higher coancestry, but here, the coancestry is measured for each pair of individuals within subpopulations. Without cross-population breeding in the experimental setup, **M** is a block diagonal matrix.

We develop a test where we estimate, in the same statistical model, the ancestral additive variances considering either (i) the variance in mean phenotype between subpopulations (*V_A,B_*) or (ii) the additive genetic variance in phenotypes within subpopulations *V_A,W_*. Under a model of neutral evolution, we expect the two estimates to be equal to the same, unique ancestral additive genetic variance. We thus test the null hypothesis that the two variances are identical:

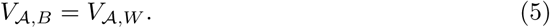

A scenario where *V_A,B_ > V_A,W_* would be compatible with local adaptation, while a scenario where *V_A,B_ < V_A,W_* would be compatible with balancing selection.

### Estimation and Framework

The use of our method requires phenotypic data of the offspring generation (which we will call F1), neutral genetic markers from the parental (𝒫) generation, and neutral genetic markers or a pedigree for the offspring generation (F1). The neutral genetic markers are used to estimate the relative mean coancestry between all pairs of subpopulations and the relatedness between all pairs of F1 individuals within subpopulations. In figure 3 we illustrate at which point the different matrices are estimated.

**Fig 3.**
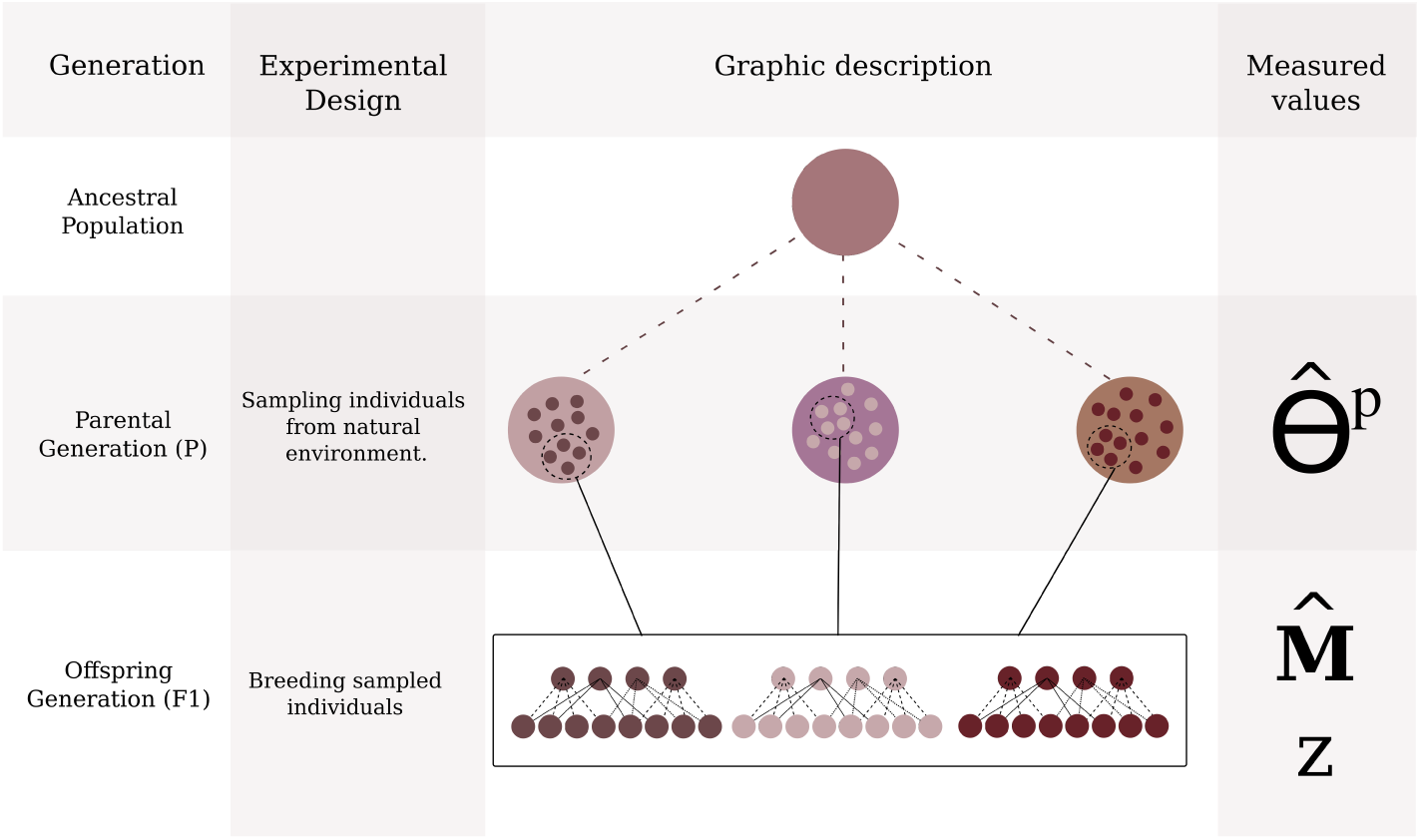
Description of expected experimental design to estimate **Θ***^p^*, **M**, and *z*. The second line of this illustrative table, the “Ancestral population” line, describes the assumed ancestral panmictic state all subpopulations derive from. The third line, “Parental generation” (𝒫) describes the studied subpopulations. From these studied subpopulations, individuals are sampled and exposed to a breeding design. At this step **Θ***^p^* is estimated. On the fourth line, we show the breeding design in the common garden setting used in our simulations: all females are crossed with all males (North Carolina II design) and two offspring from each cross are phenotyped. We estimate **M** using genotypic (or pedigree information) and phenotypic data from the F1 generation

### Estimating coancestries using the method of moments

We use the allele-sharing method of moments developed by Weir and Goudet [2017], Goudet and Weir [2023] to estimate the co-ancestry, using the minimum allele-sharing between subpopulations as a reference point. We use the method of moments to obtain **Θ***^p^* and **M** (**M** can also be obtained using pedigree information if available). The allele-sharing-based estimator can be used for species with any ploidy level and makes no assumptions about the mating system [Goudet and Weir, 2023]. For a diploid bi-allelic locus *l*, allele-sharing *A^l^_i,j_* between two individuals *i* and *j* takes a value 1 if they are homozygous for the same allele, 0 if they are homozygous for different alleles, and 1*/*2 if at least one of the two individuals is heterozygous. Over all loci, 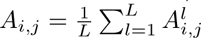.

To estimate **Θ***^p^*, we measure allele-sharing among individuals from the parental generation. The estimate 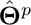 is computed as:

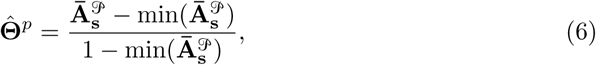

where 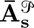 is a *r* × *r* matrix made of elements 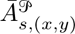, the average allele-sharing *A_i,j_* over all distinct pairs of individuals, one from subpopulation *x* and the other from subpopulation *y*; and 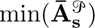 denotes the minimum off-diagonal entry of 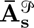 [Goudet and Weir, 2023]. This minimum entry is our best proxy for allele-sharing in the ancestral population, we therefore make the mean coancestry entries of 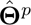 relative to the mean coancestry in the ancestral population [see Goudet and Weir, 2023, Ochoa and Storey, 2021].

Next, we need to estimate **M**, whose elements are the relatedness among the phenotyped individuals (*F*_1_) within each subpopulation discounted for their shared ancestries. To obtain relatedness, with pedigrees within each subpopulation, we could simply use the additive relationship matrix, but we can also make use of genotypic information for these *F*_1_ individuals if it is available, which we assume here. For each subpopulation, we estimate kinship among all parents and *F*_1_ individuals as

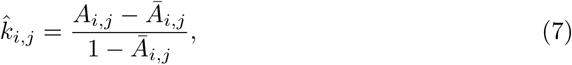

where 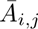 is the average allele-sharing among pairs of distinct individuals in the subpopulation [Goudet et al., 2018]. We double the kinship to obtain relatedness. To obtain 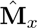, the block of the 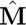 matrix corresponding to subpopulation *x*, and to account for the shared ancestry of individuals in subpopulation *x* relative to the ancestral subpopulation, we multiply the relatedness estimates of subpopulation *x* by the scalar 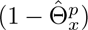, where 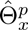 is the *x^th^* element of the diagonal of 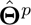. Finally we store all the 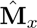 in a block diagonal matrix (see Fig. 2).

### Linear mixed model

We fit a mixed-effects model with the phenotypic trait *z* as the response variable, accounting for population and individual effects as random-effect components:

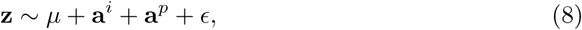

where *µ* is the overall phenotypic mean, **a***^p^* is the population-level additive genetic effect, **a***^i^* is the individual-level additive genetic effect and *ɛ* is the residual error. We assume the population-level and the individual-level additive genetic effects follow a multivariate normal distribution with their respective variance-covariance matrices depending on the above-defined co-ancestries and the between-(*V_A,B_*) and within-population (*V_A,W_*) ancestral additive genetic variances:

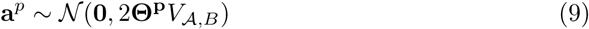

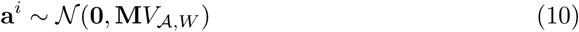

Note that assuming different ancestral additive genetic variances for **a***^i^* and **a***^p^* is another notable departure from Ovaskainen et al. [2011].

### Hypothesis Testing

We employ a Bayesian framework to estimate the ancestral additive genetic variances 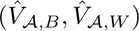 We use their posterior distributions to conduct our null hypothesis test, to incorporate uncertainty in the estimates. We test the hypothesis that the two estimates of ancestral variance are equal using the log-ratio of the ancestral variances Log*_AV_*, i.e.:

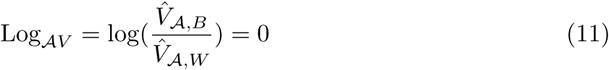

The reason for this choice is that a log-ratio is a more natural way of comparing variances than a difference, but note that associated statistical tests would be identical in both cases. From this log-ratio, we compute a Bayesian *p*-value, quantifying the proportion of the posterior sample where the sign of the log-ratio differs from the median, which has properties close to a frequentist *p*-value [Shi and Yin, 2021]. Rejecting this null hypothesis would indicate that neutral evolution alone cannot explain the observed pattern of phenotypic divergence. We refer to this method as the “log-ratio of ancestral variance” and abbreviate it as “LogAV”.

### Method testing with simulated data

We evaluated our method through simulations conducted in QuantiNemo2 [Neuenschwander et al., 2019], modeling metapopulations evolving under neutral and selective conditions. Fig 4 illustrates the population structures used for the neutrally evolving metapopulations, along with an example of a coancestry matrix following the expected pattern under neutrality. The simulations were designed to investigate population structures deviating from the standard Island Model, while also varying the number of subpopulations. We chose particularly the 1-dimensional stepping stones to give continuity to the results described in de Villemereuil et al. [2022] where the authors show the issue with the *Q_ST_* –*F_ST_* based test on a 20 subpopulation 1D stepping stones. For Stepping Stones we show results for two migration rates, leading to *F_ST_* of 0.2 and ≈ 0.37. The “139” (read: one, three, nine) population structure was inspired by the structure described by Karhunen et al. [2013] and serves as an example of a hierarchical structure. In this structure one population diverges into three subpopulations which each diverges into three new subpopulations, and the data is sampled from the nine subpopulations at the end of the simulation.

**Fig 4.**
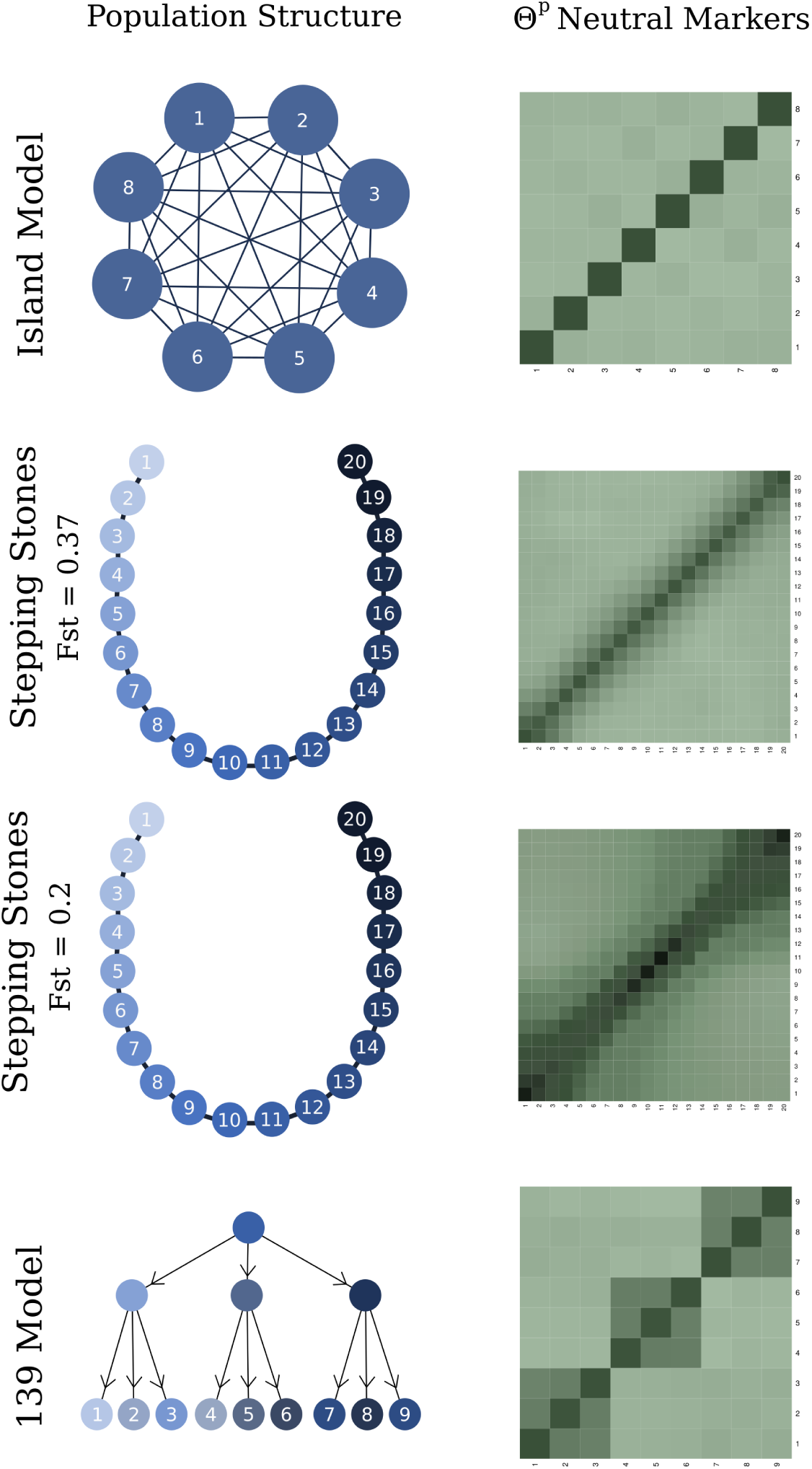
Description of studied neutrally evolving population structures along with an example of the **Θ***^p^* matrix for neutral markers.

In all simulated scenarios, we sampled ten individuals from each parental subpopulation. A common garden breeding design was implemented, wherein five females from each subpopulation were mated with five males, producing two offspring per pair. This yielded 25 families of two offspring, thus 50 offspring per subpopulation.

Phenotypes were controlled by 100 bi-allelic loci with identical and fully additive effects across all scenarios, while 2000 neutral markers were simulated for genetic analysis. In the selection scenarios, we used stabilizing selection with a different optimum between two halves of the subpopulations. In the case of the stepping stones structure, subpopulations sharing the same environment were grouped. The specific parameters for each scenario are outlined in Tables 3 and S1.

**Table 1.** Table describing relevant symbols used throughout the description of the method.

| Symbol | Definition |
| --- | --- |
| $V_A$ | Ancestral additive genetic variance. |
| $V_W$ | Additive genetic variance within subpopulations. |
| $V_B$ | Additive genetic variance between subpopulations. |
| $\hat{V}_{A,W}$ | Ancestral additive genetic variance estimated from within subpopulation effects. |
| $\hat{V}_{A,B}$ | Ancestral additive genetic variance estimated from between subpopulation effects. |
| $\Theta$ | Individual-level metapopulation-wide coancestry matrix. |
| $\Theta_x$ | Block of coancestry values related to population $x$. |
| $\Theta^p$ | Population-level metapopulation-wide coancestry matrix. |
| $\hat{\Theta}^p$ | Estimated $\Theta^p$. |
| $\Theta_x^p$ | Element $x$ of the diagonal of matrix $\Theta^p$ corresponding to coancestry within-population $x$. |
| $\mathbf{M}$ | Block-matrix of individual-level relatedness within subpopulations. |
| $\hat{\mathbf{M}}$ | Estimated $\mathbf{M}$. |
| $\mathbf{M}_x$ | A block of matrix $\mathbf{M}$ describing relatedness of individuals from subpopulation $x$. |
| $A_{i,j}$ | Allele sharing between individuals $i$ and $j$ |
| $\bar{\mathbf{A}}_s^p$ | Matrix of population-level mean allele-sharing of parental generation. |
| $z$ | Phenotypic value. |
| $\mu$ | Grand meta-population average. |
| $a^p$ | Average breeding values per subpopulation. |
| $a^i$ | Individual breeding value. |
| $\text{Log}_{AV}$ | Log-ratio of ancestral variance from between and within-population effect. |
| $n_T$ | Total number of individuals |
| $r$ | Number of subpopulations |
| $i, j$ | indices for individuals |
| $x, y$ | indices for subpopulations |

**Table 2.**
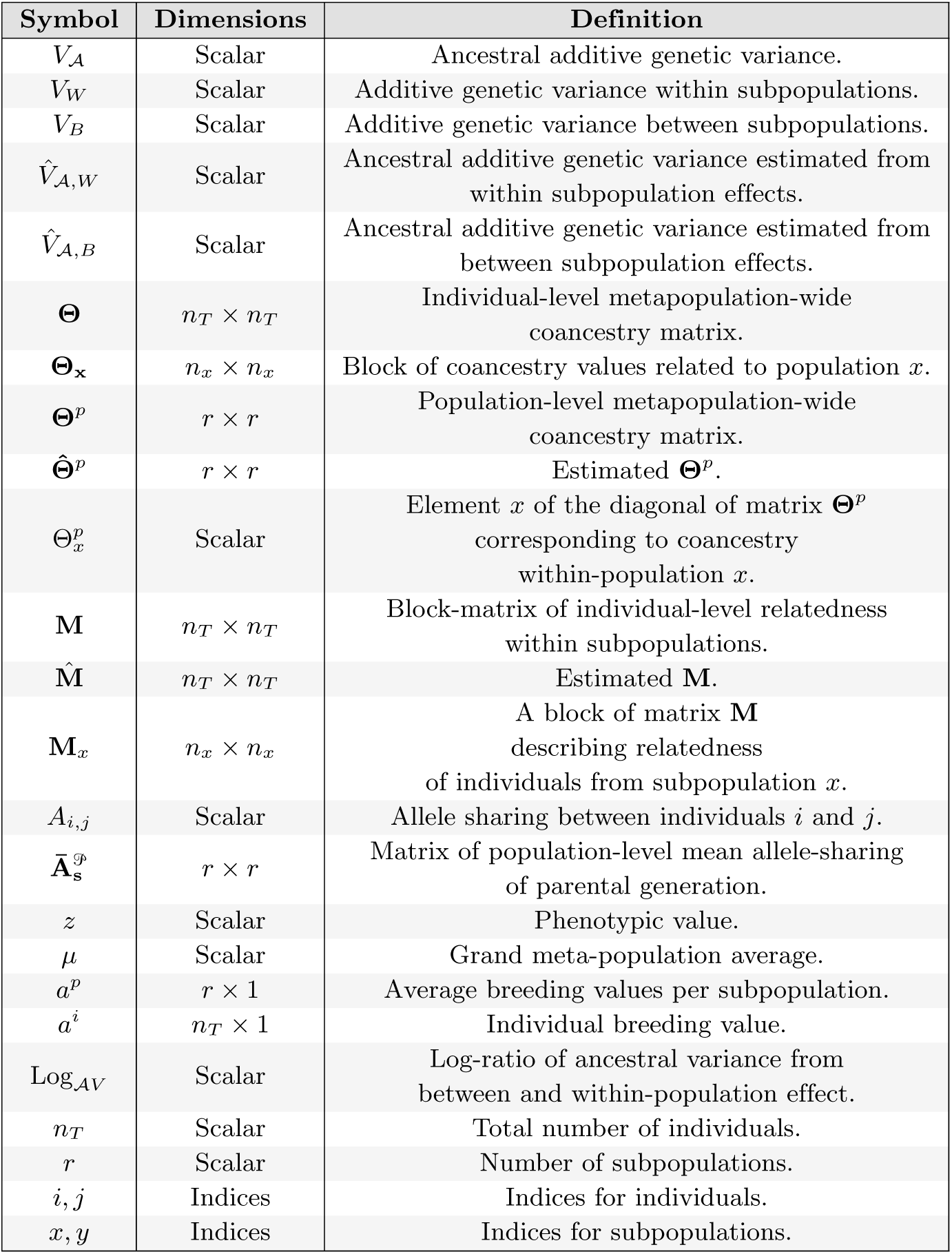
Table describing relevant symbols, their dimensions, and definitions used throughout the method.

**Table 3.**
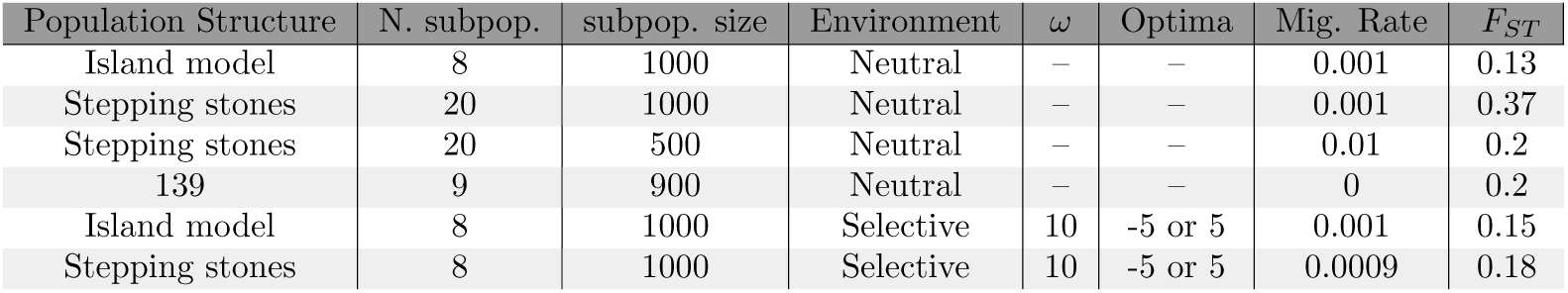
Parameters used in the different simulated scenarios.

To compare *Q_ST_* − *F_ST_* [Whitlock and Guillaume, 2009] and Driftsel Karhunen et al. [2013] to our method, we focused on the false positive rates, thus we only run these methods over neutrally evolving metapopulations. We analyzed the same set of neutrally evolving metapopulations using the three different methods, *Q_ST_* –*F_ST_*, Driftsel, and LogAV, and compared their results. By analyzing the methods over 500 replicates of each simulated neutral scenario we obtained distributions of *p*-values (in the case of *Q_ST_* –*F_ST_* and LogAV) and *S*-values (in the case of Driftsel). For our method and Driftsel Karhunen et al. [2013], we used the method of moments Weir and Goudet [2017], Goudet and Weir [2023] to estimate coancestry. We made this choice due to results in [Goudet and Weir, 2023] which show more accurate estimates of coancestry for the method of moments compared to the AFM method Karhunen and Ovaskainen [2012].

Under neutrality, *p*-values are expected to follow a uniform distribution, which is also the case for our Bayesian *p*-value [due to the asymptotical convergence of the posterior toward the sampling distribution, see Gelman et al., 2004, Shi and Yin, 2021], thus a threshold of 0.05 should correspond to a false positive rate (FPR) of 5%. The FPR is calculated as the proportion of *p*-values falling below 0.05. For Driftsel, under neutrality, *S*-values are expected to follow a normal distribution centered around 0.5 (see Supplementary Information). Following the criteria outlined by [Karhunen et al., 2013], significant *S*-values are defined as being below 0.2 or above 0.8. Although there is, strictly speaking, no theoretical expectation for what the variance of *S* should be under the null hypothesis, we choose to conform to the values advocated by [Karhunen et al., 2013] to measure a “realized” false positive rate when using their method. We performed a binomial test to compare the FPR obtained to the expected one.

## Results

Under neutrality, *p*-values obtained with the log-ratio of ancestral variance method (LogAV) are uniformly distributed for all models of population structure, including those that diverge from the Island Model, and thus yield the expected proportion of significant tests (Fig. 5). Conversely, while being correctly calibrated for the island model of population structure, the *Q_ST_* -*F_ST_* method and Driftsel show poor behavior under population structures diverging from the Island Model structure.

**Fig 5.**
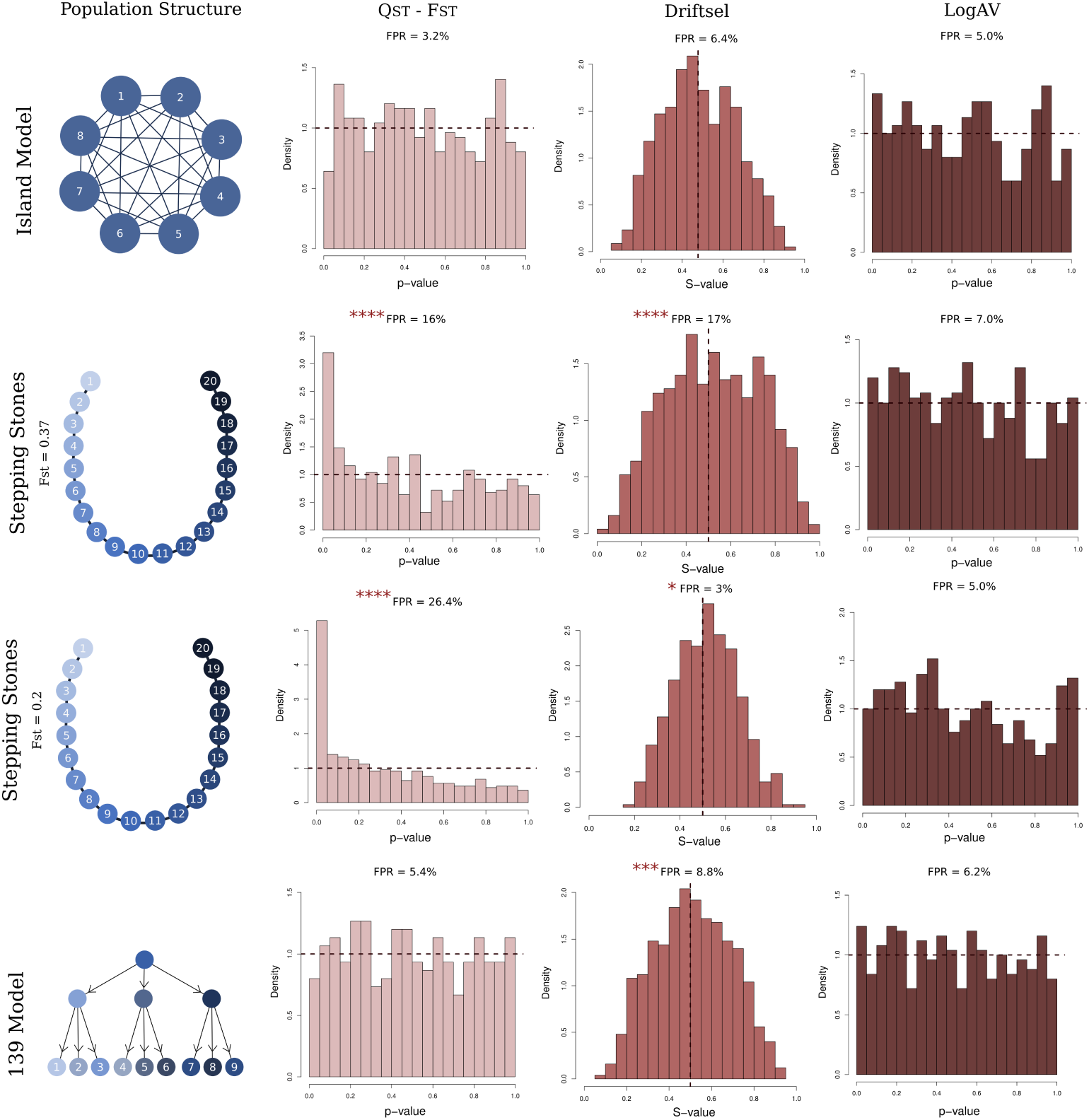
Summary of the results of analyses conducted on neutrally evolving metapopulations under varying population structures. The first column specifies the population structures examined. Subsequent columns present the outcomes for each method: *Q_ST_* –*F_ST_*, Driftsel, and LogAV. For *Q_ST_* –*F_ST_* and LogAV. The results are shown as distributions of *p*-values, while for Driftsel, the results are presented as distributions of *S*-values. False positive rates (FPR) significantly deviating from the 5% expectation are marked with asterisks.

As previously shown by [de Villemereuil et al., 2022], *Q_ST_* –*F_ST_* tests have inflated FPR under Stepping Stone population structures. We find they are significantly different 80 times out of 500 (*p*-value = 2.2 × 10*^−^*^10^) when we expect 25 significant results. Similarly, Driftsel results for Stepping Stone (*F_ST_* = 0.35) metapopulations deviate from the distribution expected under neutrality, with inflated empirical FPR (85 significant results out of 500, *p*-value *<* 2.2 × 10*^−^*^16^). Driftsel also exhibits inflated empirical FPR for the hierarchical structure 139 (44 significant results out of 500, *p*-value = 4.2 × 10*^−^*^4^). For Stepping stones with lower differentiation between demes, *i.e.* a metapopulation with average *F_ST_* of 0.2, results are qualitatively similar for *Q_ST_* –*F_ST_*. However, *Q_ST_* –*F_ST_* tests have even higher FPR. We find significance for 132 replicates out of 500 (*p*-value ¡ 2.2 × 10*^−^*^16^). As of Driftsel, we find lower FPR than the expected 5% in this lower *F_ST_* setting, with 15 out of 500. The results for neutrally evolving metapopulations are summarized in Fig. 5.

We also evaluate the LogAV test ability to detect selection. For the selection scenario we have simulated, and for the island and stepping stone models of population structures, the log-ratio of ancestral variance method shows 100% success in detecting selection, as shown in SI (Figure S2).

## Discussion

In this study, we present LogAV, a novel method based on the log-ratio of ancestral variances to disentangle the effects of selection and neutral evolution on phenotypic differentiation in spatially structured populations. This method builds on the conceptual foundation established by Ovaskainen et al. [2011], who highlighted the importance of incorporating between-population relatedness to estimate phenotypic differentiation. Through simulations, we demonstrate that LogAV is well-calibrated across all tested population structures. These simulations also showed that existing alternatives such as Driftsel and *Q_ST_* –*F_ST_*, are poorly calibrated under scenarios that assume neutrality and depart from the island model. LogAV’s reliability comes from *(i)* using coancestry matrices computed via the method of moments [Weir and Goudet, 2017, Goudet and Weir, 2023], which accurately capture genealogical ties within and between subpopulations; and *(ii)* using a theoretically justified null hypothesis test framework by comparing two estimates of the ancestral additive genetic variance. This robustness makes it well-suited for real-world use.

Real-world metapopulations often deviate significantly from the assumptions of the island model. For instance, [von Takach et al., 2022] documented a metapopulation of quolls (*Dasyurus hallucatus*) in northern Australia characterized by a hierarchical structure, comprising three main clusters with further subclusters nested within each. Similarly, the bat *Miniopterus schreibersii* demonstrates isolation-by-distance patterns, with additional fine-scale structuring observed within localities Dufresnes et al. [2023].

A good example of how structure can potentially impact studies of local adaptation is found in freshwater snails *Galba truncatula*. In western Switzerland, these snails were found in a hierarchical population structure. Trouve et al. [2005] found even physically close subpopulations, a few meters apart, showed large differentiation, and the differentiation increased as the distances between subpopulations grew. However, the geographic distance alone was not enough to describe the neutral genetic distance, with some far-away subpopulations being undifferentiated, while a small stream was sufficient to create genetic differentiation between physically close subpopulations. Using the same subpopulations, Chapuis et al. [2007] found *Q_ST_* to be significantly larger (on average) than *F_ST_* between temporary and permanent habitats, but significantly smaller among subpopulations within habitats. It is possible, however, that these results are due to not accounting for the complex population structure in this system.

Studies on local adaptation in a wide range of species have demonstrated that ecological factors strongly influence population structure. For instance, in Daphnia species, differentiation, and genetic diversity strongly depend on habitat size and type [Haag et al., 2006, Vanoverbeke et al., 2007, Walser and Haag, 2012]. Consequently, assuming the Island Model for these species may be an oversimplification, which can lead to high false positive rates in tests of local adaptation. Beyond ecology and life history traits, neutral genetic structure in metapopulations can also be shaped by historical events, such as the recolonization of previously isolated populations following glacial cycles or anthropogenic activities, as is documented for instance in the metapopulations of sticklebacks in western North America [Catchen et al., 2013]. These examples underscore the complexity of natural metapopulation structures, where migration rates and genetic connectivity often vary between pairs of subpopulations. Analyzing phenotypic divergence in such systems requires methods capable of accounting for this structural complexity. Our results show that LogAV is robust to varying population structures and should be a reliable method for testing for local adaptation in the aforementioned species.

In addition to its good performance under neutrality, the preliminary tests of LogAV using traits under selection are promising. So far, we have observed that LogAV demonstrates high sensitivity to selective scenarios. These results were consistent throughout the two different population structures tested under selection. However, further and more thorough tests, especially considering different selection strengths and sample size, are needed to identify its limits.

LogAV should be tested in scenarios that explore the interplay of various parameters, such as migration rates between subpopulations and selection patterns. Tests should include varying both the types of selection patterns and the levels of relatedness, with, for example, closely related populations being exposed to similar selective pressures. It would also be important to assess the method’s performance under varying effective population sizes, particularly in scenarios where some populations have much smaller effective sizes than others. We expect LogAV to be robust to such variations, as the inclusion of population coancestries should capture these differences in effective sizes. Additionally, the impact of sampling effort should be examined: for instance, determining the minimum number of phenotyped individuals and the number of subpopulations. Variations in relatedness within subpopulations due to the breeding systems chosen for the common garden experiments should also be explored to gauge LogAV’s robustness.

We also suggest further research to test different scenarios of ancestry. The efficacy of the proposed method depends on accurately estimating the ancestral variance from inferred relatedness, which may be impacted by assumptions of panmixia in the ancestral population. Deviations from this assumption, such as in cases of historical population substructure or admixture events, could bias variance estimates. Substructuring of the ancestral population can lead to incomplete lineage sorting and thus to non-uniform coalescent times, which could impact the estimation of the ancestral additive genetic variance. Future research should investigate to what extent the proposed method is affected by the lack of panmixia in the ancestral population.

It is important to mention that other methods have also considered the problem of population structure, and have provided potential solutions considering that little to no information on population of origin and ancestral populations is known. Josephs et al. [2019] proposed a method for detecting local adaptation using principal component analysis of the relatedness matrix. An advantage of LogAV is the more classic statistical test using an analogue of the *p*-value. Josephs et al. [2019]’s method depends on the user choosing the threshold of their tests, more specifically the cutoff for which PCs represent between- and within-population variance. In that sense LogAV is less subjective.

Further, we recognize the relevance of conducting additional research to investigate the effect of genetic architecture on the performance of methods for detecting local adaptation. Liu and Edge [2024] conducted a study that partly addresses this issue by analyzing the impact of genetic architecture on different variations of the *Q_ST_* –*F_ST_*. Specifically, they tested the impact of varying numbers of loci impacting the trait on the distribution of *Q_ST_*. Despite the theoretical expectation of *Q_ST_* not depending on the number of loci underlying the phenotype in which the statistic is measured, Liu and Edge [2024] observed that a higher number of loci leads to higher variance in the *Q_ST_* distribution. We have not explored the impact of number of loci, and it is also unclear how dominance and epistatic interactions might impact the results of our method.

Evidence suggests that the classical *Q_ST_* − *F_ST_* would be uncalibrated when there is dominance. However, the error direction is not resolved. Goudet and Martin [2007] and Cubry et al. [2017] show that dominance reduces the mean value of *Q_ST_*, meaning that dominance makes the *Q_ST_* − *F_ST_* comparison more conservative for divergent selection. While Lopez-Fanjul et al. [2007] show that under different population structures, *Q_ST_* may exceed *F_ST_*. Since the empirical effect of dominance on genetic variance needs further investigation as some studies show little influence (*e.g.* Hill et al. [2008]) of dominance while others show large influence (*e.g.* Class and Brommer [2020]), it is unclear which effects dominance would have on LogAV. Thus, it is not clear what impact dominance could have on our method, which highlights the need for future research to investigate the extent to which our method could be affected by non-additive effects on the studied phenotype.

A noteworthy difference between our method and *Q_ST_* –*F_ST_* is that it is very general regarding the breeding design used within the common garden experiment. In particular, the *Q_ST_* –*F_ST_* improvements proposed by Whitlock and Guillaume [2009] can be difficult to extend to accommodate breeding designs other than half-sib families. Specifically, their approach is constrained to balanced datasets, wherein offspring are related as half-sibs through shared fathers. Although Gilbert and Whitlock [2015] suggested an extension to unbalanced datasets, further extensions will be restricted to specific cross or nested breeding designs. Similarly to Ovaskainen et al. [2011], our method adopts a framework akin to the animal model, which creates flexibility for the breeding design. Further, and extending the flexibility of Driftsel [Karhunen et al., 2013], the within-population relatedness matrices can be obtained either from pedigree information or, alternatively, from genetic marker data in the F1 generation, enabling accurate estimation of relatedness [Goudet et al., 2018] even when the exact crosses to generate the F1 are unknown, a situation often encountered for investigations in plants, where seeds are sampled from the field and grown in a common garden environment.

In summary, our results validate the log-ratio of ancestral variance as a robust method for distinguishing adaptive divergence from neutral processes in structured populations. Its well-calibrated performance across diverse population structures demonstrates that it provides a reliable alternative to *Q_ST_* –*F_ST_* and Driftsel. We did so by adopting a general framework based on well-established quantitative genetic and statistical principles. From quantitative genetic theory, we derive two distinct approaches to calculate ancestral additive genetic variances, *V_A_, W* and *V_A_, B*. Then we compare the two estimates of ancestral variance using a log-ratio approach that has been adopted by many hypothesis testing frameworks (e.g., likelihood ratio test) and has proven to be very robust in a wide range of applications requiring the comparison of variances.

## Acknowledgments

We thank Otso Ovaskainen and Juha Merila for fruitful discussions and suggestions.

## Supporting information

### *S*-statistics results

#### The ***S***-statistic and its behavior under neutrality

The *S*-statistic is a measure proposed by [Ovaskainen et al., 2011] to assess deviations from neutral expectations in population-level effects of quantitative traits. It quantifies how likely the observed pattern of population differentiation described by the estimated population-level random effects (**a***^p^*) given the expected neutral distribution of population-level effects obtained from random realization of the distribution(**a***^p^_R_*). The neutral expectation is modeled as a multivariate normal distribution with covariance structure dependent on the metapopulation-level coancestry matrix (**Θ***^p^*). Then to compute *S*, we compare the probability density of the observed population-level effects under the neutral expectation to the probability density of random realizations of the neutral model.

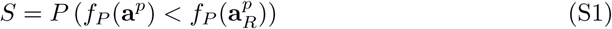

An *S*-value close to 1 indicates that the observed pattern is highly unlikely under neutrality, signaling potential selection, whereas *S* ≃ 0.5 suggests a pattern consistent with neutral expectations. *S* ≃ 0, indicates that the observed population effects are more consistent than expected under neutrality. Under neutrality, the distribution of *S* is symmetric around 0.5 because *f_P_* (*a^P^*) and *f_P_* (*a^P^_R_*) are equally likely to be greater or less than each other. This symmetry arises from the inherent properties of the neutral model, where population-level effects follow a multivariate normal distribution centered on the ancestral mean. Without selection, deviations in either direction (larger or smaller likelihoods for *f_P_* (*a^P^*) compared to *f_P_* (*a^P^_R_*)) occur with equal probability. However, the exact distribution of *S* under neutrality is not obvious.

#### Asymptotic normality of the posterior mean of ***S***

Driftsel Karhunen et al. [2013] estimates *S* using a Bayesian method that fits the model parameters and generates posterior samples. Since we are using independent replicates of simulated data, each sample provides a different estimate of *S*, and the mean S-statistic is calculated as the average across the posterior samples. Thus the average of a sufficiently large number of independent and identically distributed random variables approaches a normal distribution, regardless of the underlying distribution of individual variables.

### Computation of the ***p***-value associated with the LogAV

Given the posterior distribution of the log-ratio of ancestral variances *P* (Log*_AV_* |*D*) (calling *D* the total set of data, including information regarding the traits and the co-ancestries), we compute the two-tailed Bayesian *p*-value associated with the null hypothesis that Log*_AV_* = 0 as:

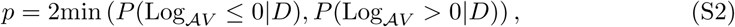

where the factor 2 is a scaling factor arising from accounting for the conditional probability of picking one of the probabilities in the minimum function (which happens to be 0.5 under the null hypothesis). Given the asymptotic property that posterior distributions reflect long-term sampling distribution of the estimators, with sufficiently weakly informative priors [Gelman et al., 2004], such constructed *p*-values tend to behave like frequentist *p*-values, and especially, to be uniformly distributed [Shi and Yin, 2021]. This is tightly connected to the asymptotic behavior of credible intervals as confidence intervals under similar assumptions [Gelman et al., 2004]. Although long-term frequentist interpretation of such *p*-values (and null hypothesis testing framework in general) is not the most natural use of Bayesian statistics [Gelman et al., 2004], we chose to use it here because *(i)* such *p*-values are easily obtained from a particular model compared to alternatives like information criterium based model selection; *(ii)* it tracks back with habits established in the field to use null hypothesis testing for *Q_ST_* –*F_ST_* comparison; and most importantly, *(iii)* the biological question tackled here fundamentally requires to test the null hypothesis of neutral evolution.

### Comparison of LogAV and ***Q_ST_*** –***F_ST_*** results

Here we compare the distribution of *p*-values between LogAV and *Q_ST_* –*F_ST_*Whitlock and Guillaume [2009] using the same sampled individuals and measured phenotypes.

### Selection results

#### Simulation parameters per scenario

We simulated different demographic and selective scenarios, each run with 500 replicates. All models assumed biallelic loci for both neutral and quantitative trait loci, a mutation rate of *µ* = 10*^−^*^7^, and freely recombining loci. The trait heritability was *h*^2^ = 0.8. Detailed simulation parameters can be found in S1.

#### Supplementary Materials: Simulation Parameters

**Table S1.** Summary of simulation parameters for each scenario.

| Scenario | Generations | Subpop. Size | Num. Subpop. | Sample per Subpop. |
| --- | --- | --- | --- | --- |
| Neutral IM | 500 | 1000 | 8 | 10 |
| Neutral SS low Fst | 5000 | 500 | 20 | 10 |
| Neutral SS high Fst | 5000 | 1000 | 20 | 10 |
| Neutral 139 | 900 | 1000 | 9 | 10 |
| IM selection | 500 | 1000 | 8 | 10 |
| SS selection | 500 | 1000 | 8 | 10 |

The selection equation applied in the island and stepping stone models under selection was:

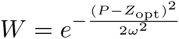

where *W* is fitness, *P* is the trait value, *Z*_opt_ is the optimal trait value (5 for demes 1–4 and −5 for demes 5–8), and *ω* = 10 represents the strength of stabilizing selection.

### Method testing results compared to theoretical expectation

On figures S3, S4, and S5 we show qq-plots displaying the difference between the theoretical distribution shapes and the observed ones for the different methods. While the *S*-statistics neutral distribution for the many replicates should follow a normal distribution, the expectation is uniform for the *p*-value distribution of the other two methods.

---

**Fig S1.**
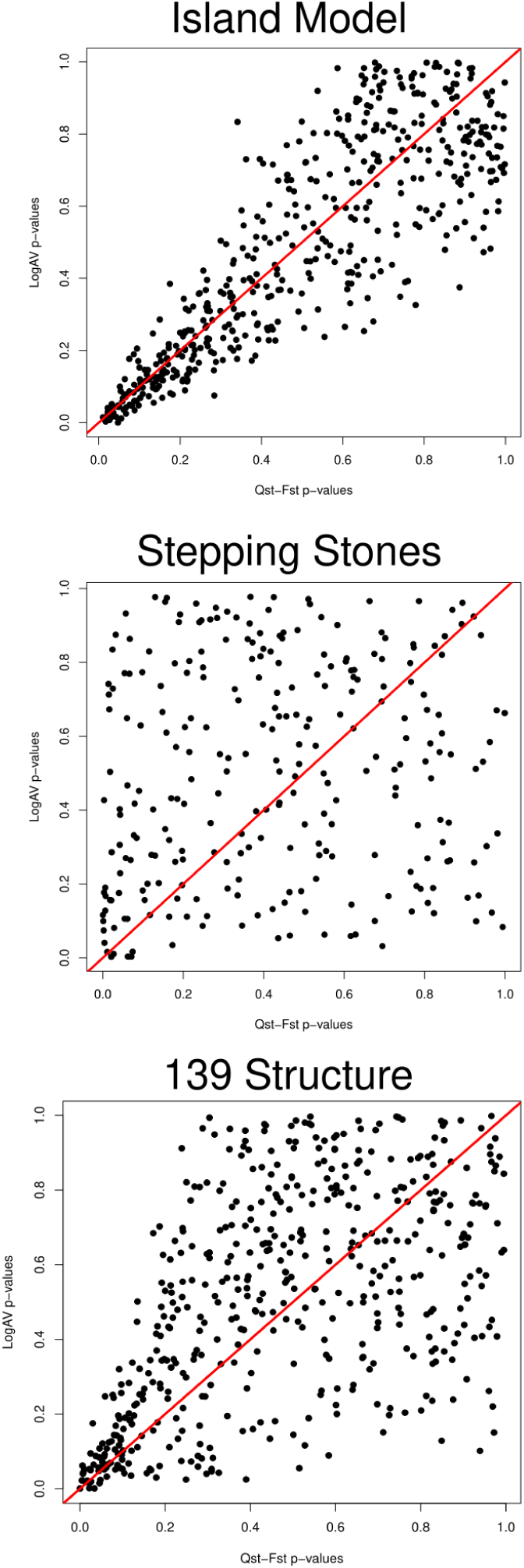
Comparison between distributions of results of the LogAV method and *Q_ST_* –*F_ST_* Whitlock and Guillaume [2009] over neutraly evolving simulated metapopulations under Island model, Stepping stones, and 139 structure

**Fig S2.**
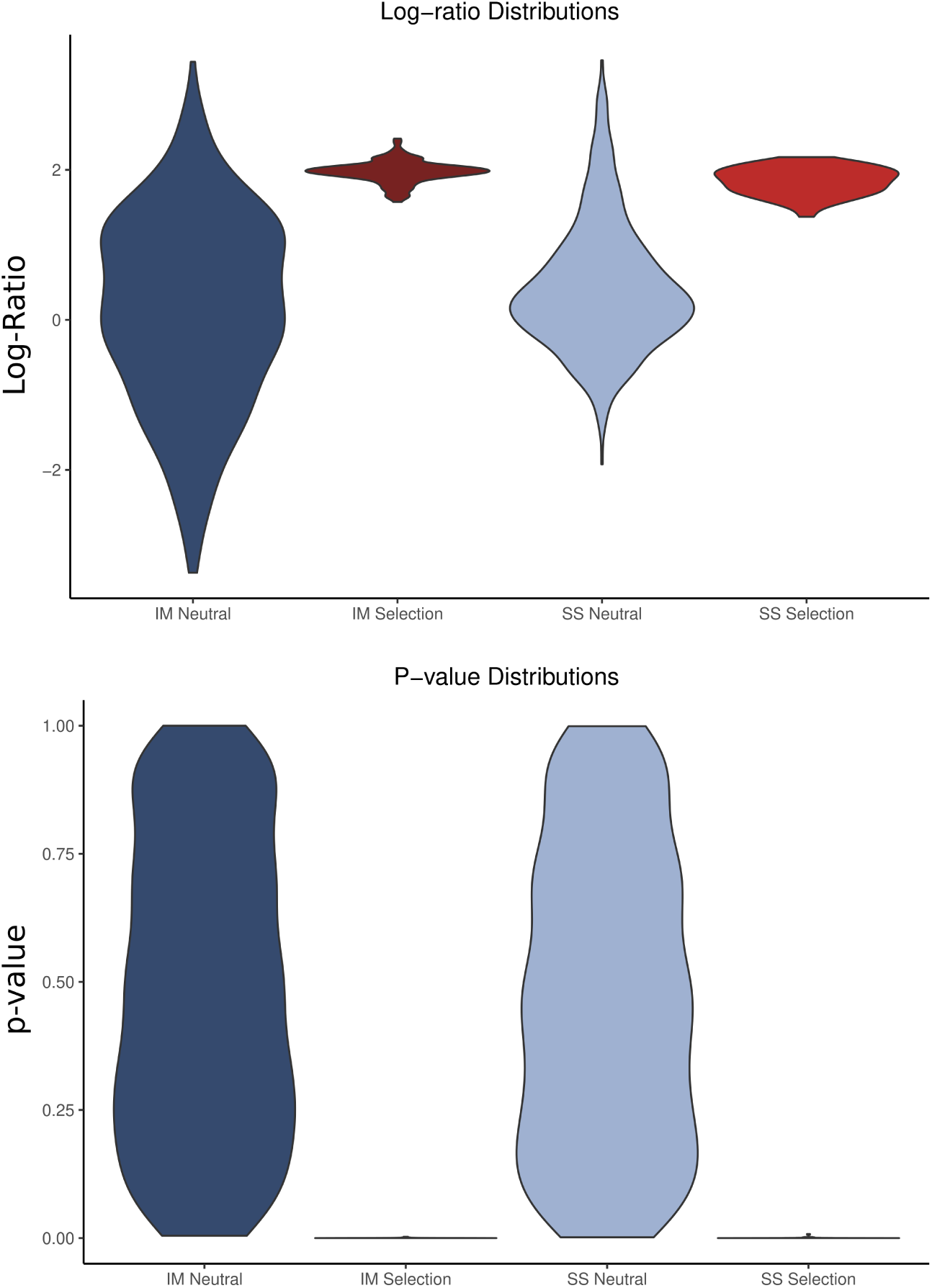
Comparison between distributions of results for Island model under neutrality, Island model under selection, Stepping stones under neutrality and Stepping stones under selection. The top figure shows the distribution of the Log-Ratio of 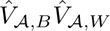, while the bottom figure shows the *p*-value distribution.

**Fig S3.**
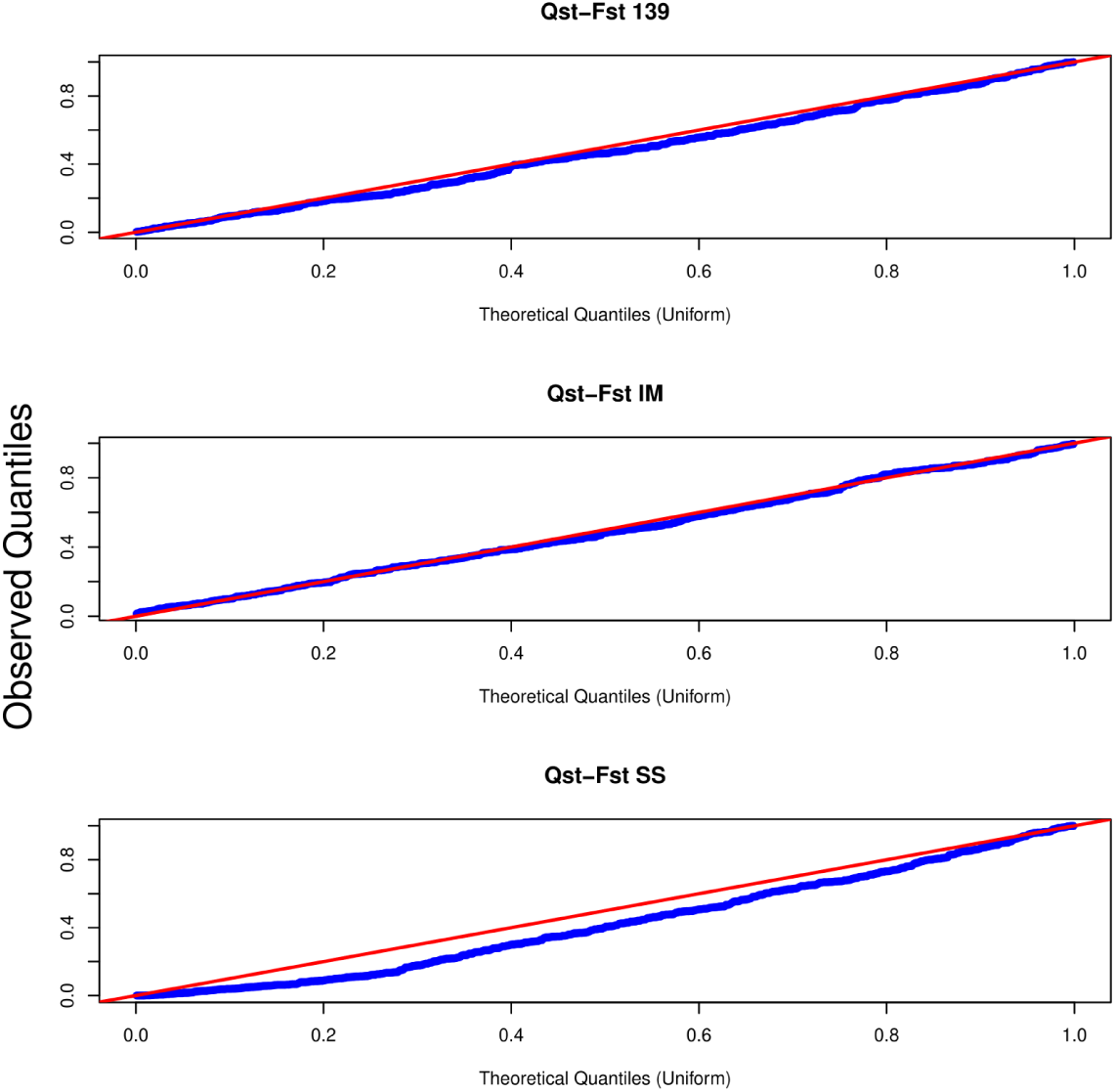
Quantile plots comparing the theoretical expectation under neutrality with the observed distribution for *Q_ST_* -*F_ST_*. We show the comparison for three neutrally evolving population structures.

**Fig S4.**
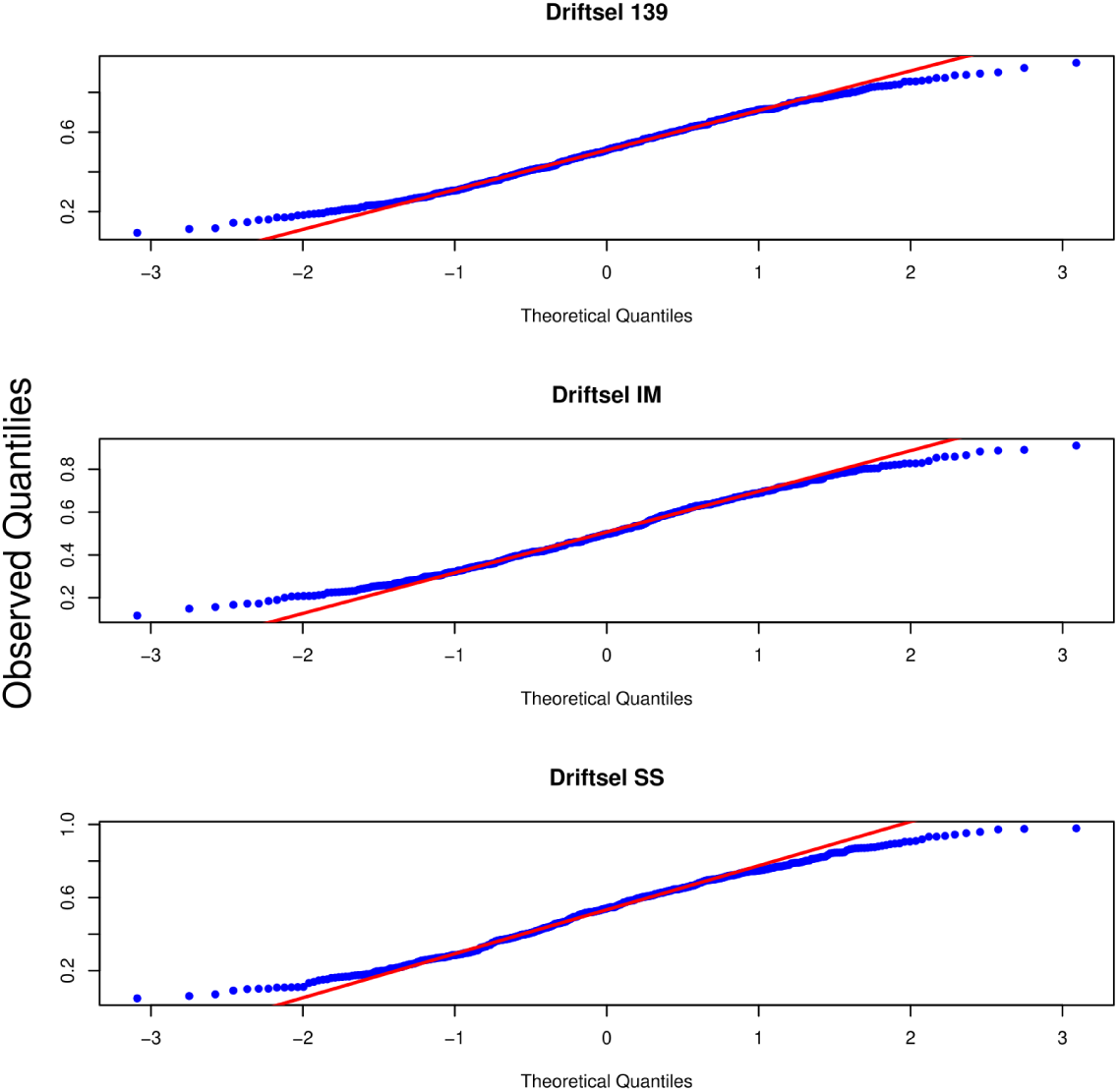
Quantile plots comparing the theoretical expectation under neutrality with the observed distribution for Driftsel. We show the comparison for three neutrally evolving population structures.

**Fig S5.**
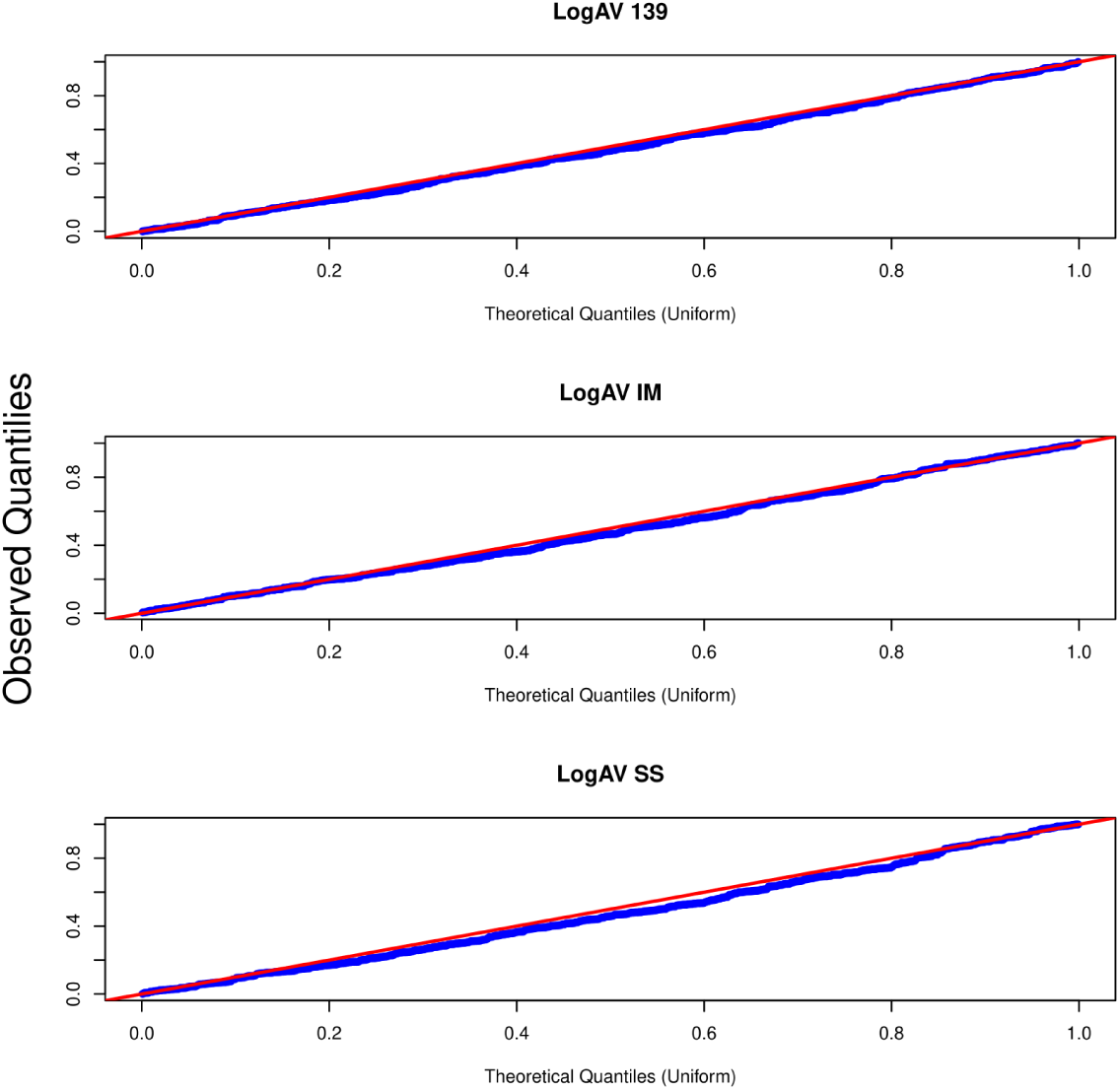
Quantile plots comparing the theoretical expectation under neutrality with the observed distribution for LogAV. We show the comparison for three neutrally evolving population structures.

